# Spatiotemporal relationships defining the adaptive gating of the bacterial mechanosensitive channel MscS

**DOI:** 10.1101/154435

**Authors:** Uğur Çetiner, Sergei Sukharev

## Abstract

Adaptive desensitization and inactivation are common properties of most ion channels and receptors. The mechanosensitive channel of small conductance MscS, which serves as a low-threshold osmolyte release valve in most bacteria, is unusual because it slowly inactivates not from the open, but from the resting state under moderate tensions. The manifestation of this mechanism is the channel’s ability to discriminate the rate of tension application, i.e., to ignore slow tension ramps but fully respond to abruptly applied stimuli. In this work, we present a reconstruction of the landscape for tension-dependent MscS transitions based on patch current kinetics recorded under specially designed pressure protocols. The data are analyzed with a three-state continuous time Markov model of gating, where the tension-dependent transition rates are governed by Arrhenius-type relations. The analysis provides assignments to the intrinsic opening, closing, inactivation, and recovery rates as well as their tension dependencies. These parameters, which define the spatial (areal) distances between the energy wells and the positions of barriers, describe the tension-dependent distribution of the channel population between the three states and quantitatively predict the experimentally observed dynamic pulse and ramp responses. Our solution also provides an analytic expression for the area of the inactivated state in terms of two experimentally accessible parameters: the tension at which inactivation probability is maximized, γ*, and the midpoint tension for activation, γ_0.5_. The analysis initially performed on *Escherichia coli* MscS shows its applicability to the previously uncharacterized MscS homolog from *Pseudomonas aeruginosa*. MscS inactivation minimizes metabolic losses during osmotic permeability response and thus contributes to the environmental fitness of bacteria.

## INTRODUCTION

Mechanosensitive (MS) ion channels are found in all domains of life. While in animals they fulfill multiple sensory functions (Alloui et al., 2006; Heurteaux et al., 2006; Retailleau and Duprat, 2014; Volkers et al., 2015; Walsh et al., 2015), in prokaryotes (Kung et al., 2010), protozoans (Prole and Taylor, 2013) and plants (Nakagawa et al., 2007, Haswell et al., 2008) they participate in tension-triggered redistribution of osmolytes to balance osmotic forces in different compartments. In enteric bacteria, which are transmitted between hosts through fresh water and thus are frequently subjected to drastic shifts of external osmolarity, the osmolyte release system is particularly robust. In order to survive an abrupt 750 mOsm downshift causing massive water influx, *E. coli* cells eject most of their small osmolytes (up to 20% of cell’s dry weight) within ~50 ms (Çetiner et al., 2017). Smaller bacteria, such as *Pseudomonas aeruginosa*, do it even faster (Çetiner et al., 2017). This massive but fully reversible osmotic permeability response is mediated primarily by two MS channels: the low-threshold MscS and the high-threshold MscL residing in the inner membrane ( Sukharev et al., 1993; Blount et al., 1996; Levina et al., 1999;). The 3-nS MscL expressed in wild-type *E. coli* strains is an emergency valve that opens at near-lytic tensions (10-14 mN/m) in the event of a strong abrupt shock (Levina et al., 1999; Sukharev et al., 1999). The 1-nS MscS, in contrast, opens at around 7.8 mN/m and shows adaptive behavior involving complete inactivation wherein it enters a non-conductive and tension-insensitive state (Akitake et al., 2005, 2007). MscS alone can effectively counteract small shocks by curbing tension before it reaches the threshold for MscL (Levina et al., 1999). At strong shocks, which open MscS and then MscL, we expect a close cooperation between the channels. When MscL opens and dissipates the bulk of osmolytes, it lowers the tension to its own threshold and then closes. At this point, MscS is still supposed to be open and to continue the release process until it reduces the membrane tension well below the MscL threshold. When the tension approaches the MscS threshold, the channel closes and finally inactivates, completely resealing the membrane. MscS inactivation appears to exclude spurious flickering of channels at “safe” near-threshold tensions, thus minimizing metabolic losses and assisting in a more complete recovery from osmotic shock.

Previous analysis has shown that the transition into the inactivated state occurs only from the closed state under moderate tensions, specifically in the range that activates only a fraction of the population and requires at least a few dozen seconds to manifest (Kamaraju et al., 2011) (See Supplementary Fig. 1). Therefore, the channel operates in two separable time scales. For a quick application of strong stimuli, the channels are distributed primarily between the open and the closed states. For slow or prolonged application of moderate stimuli that keeps the majority of channels closed, the population gradually redistributes into the non-conductive and tension-insensitive inactivated state. The opening and inactivation transitions are both originated from the closed (C) state, thus, the outcome of their competition, i.e. population fractions in the open (O) and inactivated (I) state, is determined by transition rates and their tension dependences (Fig. 1).

**Figure 1.**
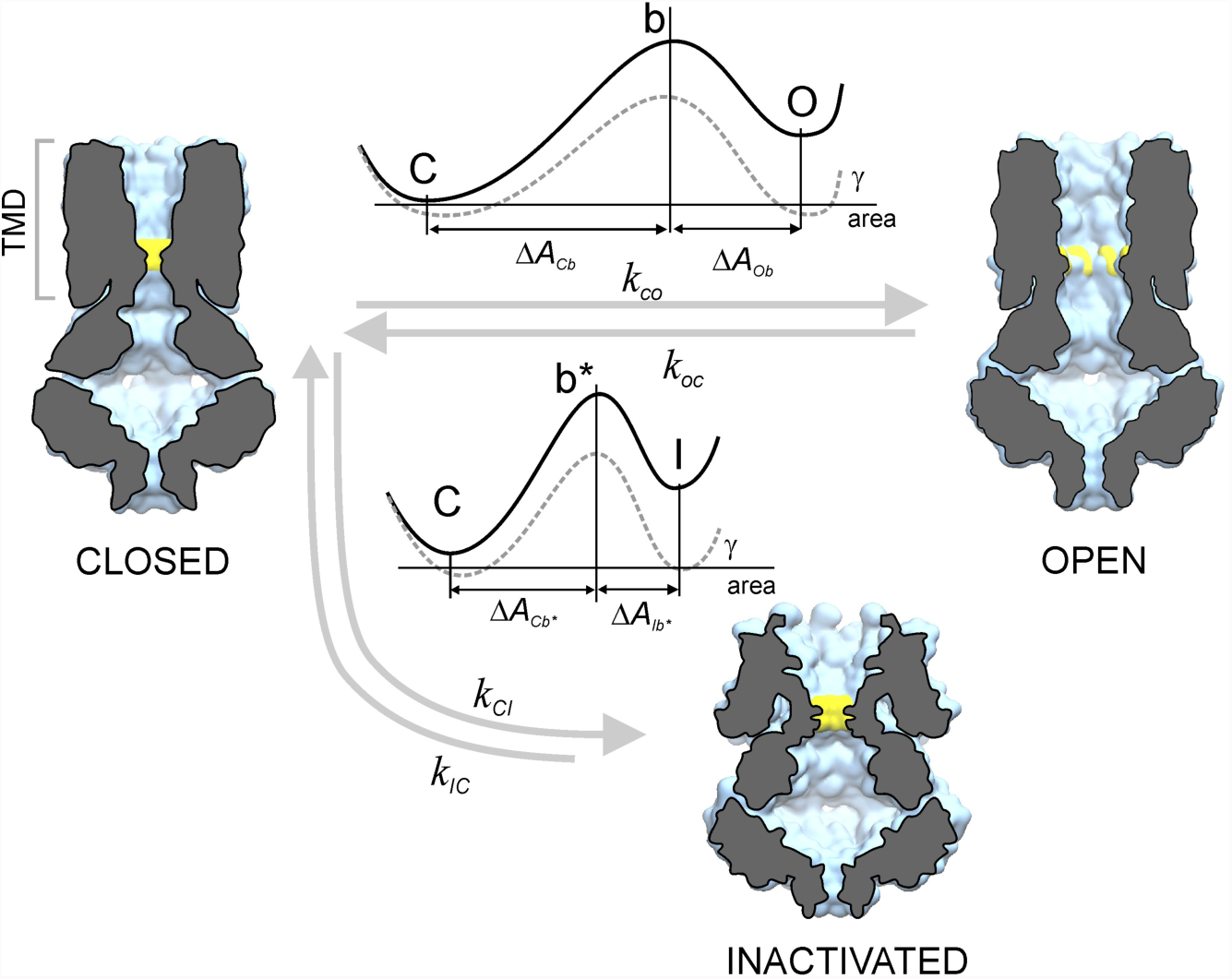
The discrete conformational state space of allowed transitions indicated by the experiments. The free energy difference between the states decreases linearly with the work done by the tension, γΔA,, favoring the states with larger in-plane area (A) of the transmembrane domain (TMD). The transition rates between the states are governed by an Arrhenius type of relation displaying exponential dependence on the membrane tension. The barriers that separate the open and inactivated states from the closed state are denoted by b and b*, respectively.

Since both the C ➔ O and C ➔ I transition rates increase with tension transmitted laterally through the lipid bilayer, each of the transitions is expected to be accompanied by a protein area increase in the plane of the membrane. The scale of area increase from the closed well to the transition barrier defines the slope of the kinetic rate vs. tension, whereas the area increase between the wells would define how the equilibrium partitioning between the wells changes with tension (Sukharev et al., 1999, Kamaraju et al., 2010). The process with the steepest rate dependence (in this case - the opening) is expected to dominate at high tensions.

Crystallographic structures, modeling and molecular dynamics simulations gave us some insight into the molecular nature of the two competing processes. In brief, the crystal structure of MscS (2OAU) solved in the absence of surrounding lipids appears to be non-conductive (Anishkin and Sukharev, 2004; Sotomayor and Schulten, 2004; Sotomayor et al., 2006; Anishkin et al., 2008a) and at the same time shows the peripheral lipid-facing helices (TM1-TM2) in a splayed conformation, uncoupled from the gate. This non-conductive, uncoupled structure was therefore interpreted as the inactivated state (Anishkin and Sukharev, 2004; Akitake et al., 2005, 2007). Re-packing of the peripheral TM1-TM2 helices in a fashion more parallel to the pore-lining TM3s produced a compacted resting (closed) state with re-formed TM2-TM3 contacts capable of transmitting tension from the bilayer to the gate. From that state, under high (near-saturating) tensions the channel can open through complete straightening of its TM3 helices (Akitake et al., 2007; Anishkin et al., 2008b). Starting from the compact resting state, the channel, therefore, can undergo alternative transitions to either the open or inactivated state, each characterized by a specific area change (Fig. 1).

In this paper, we perform experimental patch-clamp analysis of the MscS kinetics under programmed pressure protocols designed to reveal rates for specific transitions. We introduce a finite state continuous time Markov chain model, extract intrinsic rates and their tension dependencies, and then perform time-dependent kinetic simulations which reasonably reproduce experimental traces obtained with both step and ramp protocols. The model quantitatively reproduces the ability of MscS to reduce its response to slowly applied stimuli previously described as the ‘dashpot’ behavior (Akitake et al., 2005). The parameters indicate spatial relationships between the energy wells and separating barriers, thus reconstructing the tension-dependent landscape for the transitions. We show that the model based on analysis of *E. coli* MscS with interconnected parameters can be applied to MscS homologs from other organisms, which are presently characterized in less detail.

## Methods

### Theoretical model development

The mathematical model consists of a discrete-state continuous time Markov chain where the transition rates are described by the Arrhenius relation and the dynamics satisfy the detailed balance condition. For instance, the transition rates for the closed ↔ open transition obey the following relation: 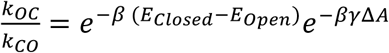. Thus, in the presence of membrane tension, not only do the states with larger areas become more favorable but also the detailed balance condition guarantees that once the tension is held fixed the system relaxes to a unique equilibrium distribution where the states are populated according to Boltzmann weight (Ritort, 2004). The three-state model does not preclude the existence of some short-lived sub-conducting or hidden Markov states. Its applicability will be tested in the experiments described below. For convenience of relating tension with the energetic bias toward lateral expansion, the membrane tension here is mostly presented in units of k_B_T/nm^2^, which equals to 4.114 mN/m at room temperature (298K).

### Experimental electrophysiological recordings and data treatment

The WT *E.coli* MscS was the primary object for patch-clamp recording. Additionally, the *P. aeruginosa* MscS-1 channel, the recently cloned close homolog of *E. coli* MscS (Çetiner et al., 2017), was used for comparison. Both MscS homologs were expressed from the pB10d vector under the PlacUV5 inducible promoter (Okada et al., 2002) in the MJF641 strain devoid of seven endogenous MS channel genes (Δ*mscL, mscS, mscK, ybdG, ynaI, ybiO and yjeP*) (Edwards et al., 2012). Giant *E. coli* spheroplasts heterologously expressing the channels were generated with a standard method as described previously (Akitake et al., 2005; Martinac et al., 1987).

Borosilicate glass (Drummond 2-000-100) pipets 1-1.3 *μ*m in diameter were used to form tight seals with the inner membrane. The MS channel activities were recorded via inside-out excised patch clamp method after expressing them in MJF461. The pipette solution had 200 mM KCI, 50 mM MgCl_2_, 5 mM CaCl_2_, 5 mM HEPES, pH of 7.4. The Bath solution was the same as the pipette solution with 400 mM Sucrose added. Traces were recorded using the Clampex 10.3 software (MDS Analytical Technologies). Mechanical stimuli were delivered using a modified high-speed pressure clamp apparatus (HSPC-1; ALA Scientific Instruments). Current data were analyzed after Rs correction of the traces by the equation, Gp=I/ (V-IRs), where Gp is the patch conductance and V and I are the transmembrane voltage and current, respectively. Rs is the intrinsic resistance of the patch free pipette (1.2-2 MΩ). The pressure (P) was converted to tension (*γ*) using the following relation: *γ* = (*P*/*P*_0_._5_)*γ*_0.5_ assuming the radius of curvature of the patch does not change in the range of pressures where the channels were active and the constant of proportionality between tension and pressure was taken to be *γ*_0.5_/*P*_0.5_ (Sukharev et al., 1999). The midpoint tension *γ*_0.5_ of MscS was taken to be 7.85 mN/m (Belyy et al., 2010a) and the midpoint tension of PaMscS-1 activation in *E. coli* spheroplasts was determined to be 5.6 mN/m using MscL as an intrinsic tension gauge (Çetiner et al., 2017). *P*_0.5_ values that correspond to pressure at which half of the population is in the open state were determined using 1-s triangular ramp protocol.

## Results

In this section, we will first introduce the model and obtain some analytical results for the steady state behavior. Next, we will present the main experimental observations followed by the thermodynamic and kinetic treatment of the system. Following this, we will summarize the kinetic and spatial parameters extracted from the data and then finally simulate the responses and compare them with experimental traces to confirm the quality of the model. We should note that the experimental protocols were developed previously (Akitake et al., 2005; Kamaraju and Sukharev, 2008; Belyy et al., 2010; Kamaraju et al., 2010, 2011), but the datasets presented in previous papers on MscS produced only partial quantitative information and have never been treated comprehensively.

### The three-state continuous time Markov chain model

Fig. 1 presents the kinetic scheme of the MscS functional cycle. The unperturbed channel resides in its native environment in the compact closed (resting) state. From that state, the channel undergoes two separate tension-driven transitions into the open (C ➔ O) or inactivated (C ➔ I) states. Note that the O and I sates are not interconnected, meaning that open channels do not inactivate (Kamaraju et al., 2011), (See Supplementary Fig 2). The energy wells for the connected states are assumed to be separated by single rate-limiting barriers and the transition rates between the states are governed by the Arrhenius-type relation: 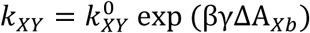 where 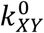 is the intrinsic rate (frequency) of the system’s attempts to overcome the barrier between states X and Y in the absence of tension (Bell, 1978). The exponential term includes Δ*A*_xb_, the expansion area from the bottom of the well of the particular state, X, to the top of the barrier that separates the states X and Y, γ, is the applied tension, and β=1/k_B_T. Thus, tension favors states with larger area.

**Figure 2.**
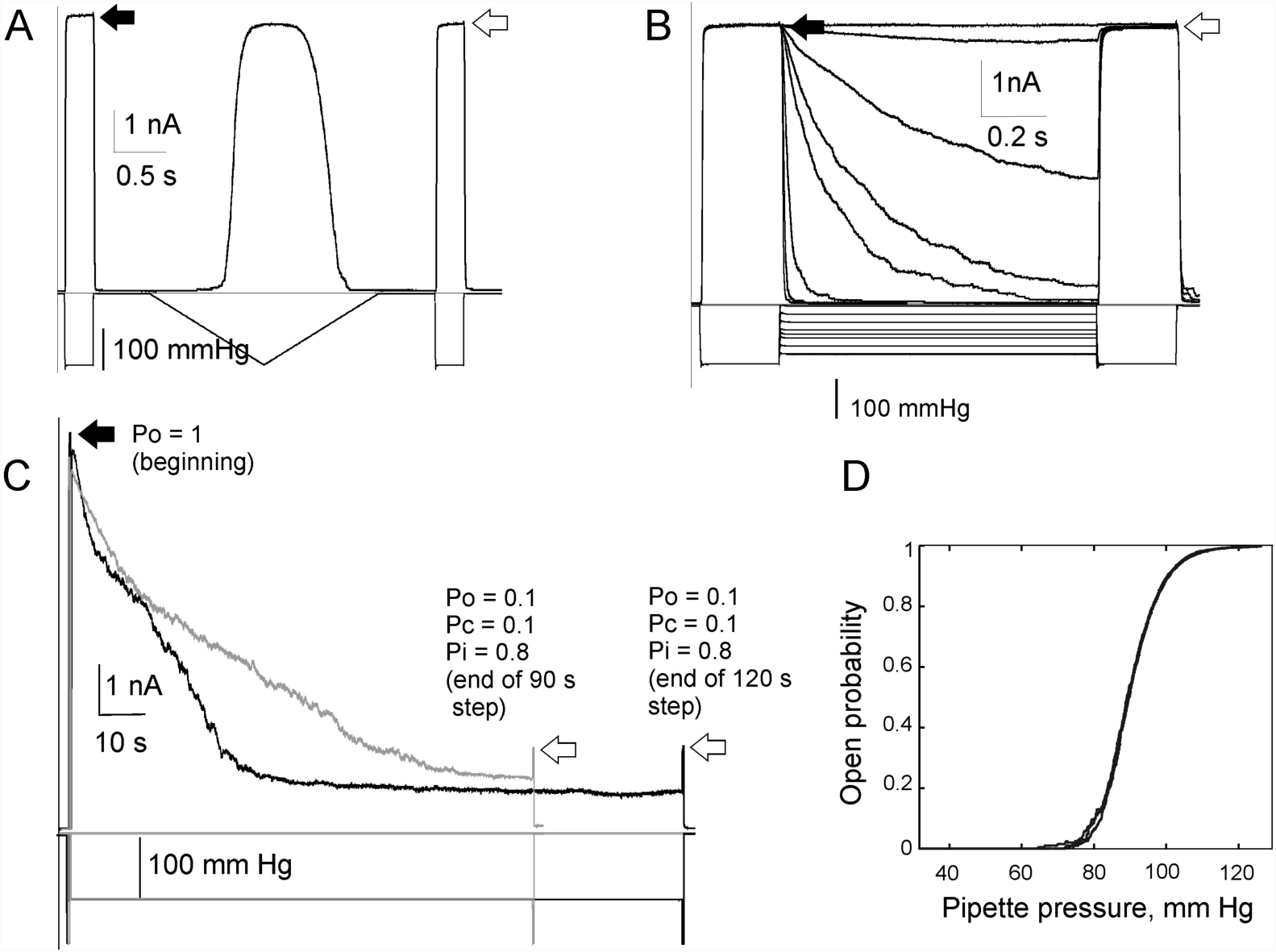
Exploration of time scales characteristic for the separate C↔O and C↔I transitions in MscS populations. (A) Current responses to a short pulse-ramp-pulse stimulus. (B) Responses to a series of short (1s) pulse-step-pulse stimuli. Application of quick ramp or pulse protocols does not induce any inactivation thus MscS can be well modeled as a two state system. (C) Superimposed 90s and 120 s pulse-step-pulse experiments showing that a 90 s step is sufficient for reaching equilibrium between the C, O and I states. (D) Control ramp experiments before, in between, and after 90 and 120s experiments illustrating a good superimposition of three activation curves and intactness of the patch excluding slippage inside the pipette. The black and white arrows indicate maximal amplitudes of population current before and after the ramp or step stimulation.

In this kinetic framework, when the tension is kept constant, the master equation can be written as follows:
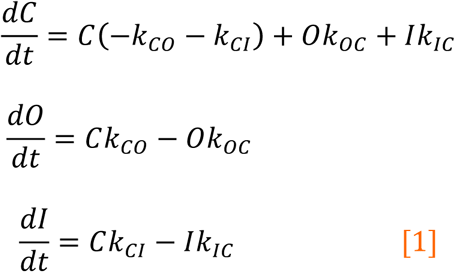

Or in a more compact form:
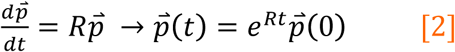

where 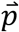 is the probability vector and R is the transition rate matrix specified as:
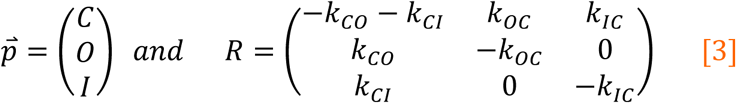

The equilibrium distribution of the chain can be obtained in several ways. Perhaps the simplest is to set 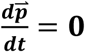 as a stationary solution and solve a system of linear equations. Another way is to use the exponential form for the probability vector (Schnakenberg, 1976; Steinfeld et al., 1989), which can be written as:

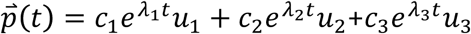

Where *λ*_1_, *λ*_2_, *λ*_3_ are the distinct eigenvalues of R and *u*_1_, *u*_2_, *u*_3_ are the corresponding right eigenvectors. The Perron-Frobenius theorem then guarantees that there is going to be a unique invariant distribution, ***π***, that can be expressed as: 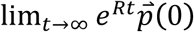. A detailed discussion on Perron-Frobenius theorem for primitive matrices, convergence and uniqueness for finite state, and irreducible Markov chains is included in the supplementary information (Norris, 1998).

The equilibrium distribution of the inactivated state probability is given by the following equation:
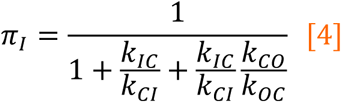

By plugging in the relations for the transition rates, we obtain:
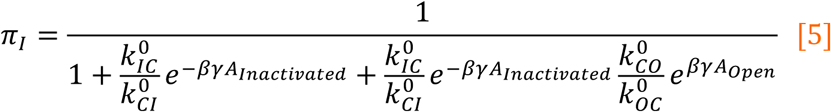

where *A_Inactivated_* = Δ*A*_*Ib*\*_ + Δ*A_cb_*_*_ and *A_open_* = Δ*A_cb_* + Δ*A_ob_* represent the total in-plane area expansion associated with the inactivated and the open state respectively, assuming there is a single barrier separating the states. Since low-tension values favor the closed state and high-tension values lock the system in the open state with no transition to the inactivated state, we assume there is an optimum intermediate tension that simultaneously allows for a sufficient fraction of channels to be left in the closed state and yet also provides enough driving force to maximize the probability of inactivation. In the following analysis, we determine the tension *γ*\* that gives the maximum for inactivated state probability by solving the system under the 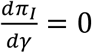 condition. As detailed in the supplement, the exact expression for the tension at which the probability of inactivation reaches its maximum is given by:
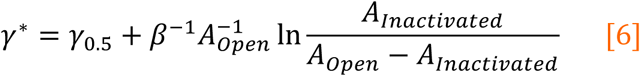

where *γ*_0.5_ is the tension at which probability of finding a channel in the open state is 0.5. This expression, therefore, relates the tension that results in the highest degree of inactivation with the half-activating tension. This expression can be rewritten in terms of the inactivated state area, *A_Inactivated_*, which reflects the in-plane expansion of the channel protein associated with the inactivation transition,
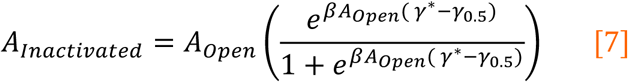

As was reported previously, *A_Inactivated_* is of prime mechanistic interest because it reflects the de-coupling of the peripheral helices from the gate (Akitake et al., 2007; Belyy et al., 2010b). However, it is not an easily accessible parameter(Kamaraju et al., 2011). Having this relationship we now notice that *A_open_*, *γ*\*, *γ*_0.5_ are easier to obtain, thus the area of the inactivated state can be found using the expression [7] without a need for more experimental parameters.

## Experimental Results

### 1. The time scales for C↔O and C↔I transitions

As shown on the kinetic scheme (Fig. 1), the transitions between C and O and C and I states are governed by different barriers and therefore the equilibration upon stimulus application may take different times. We adopted the previous strategy of recording the transition with either ramps or prolonged steps while controlling the active fraction of the population with short saturating test pulses, before, after, and in some cases during extended stimulation. Fig. 2 shows MscS population responses under stimuli applied in two different time scales. Panel A shows a response to a pulse-ramp-pulse sequence, where 0.2 s saturating test pulses flank a 2-s symmetric ramp (1-s ascending and 1-s descending limbs). The bell-shaped ramp response is slightly asymmetric showing slight hysteresis at this ramp rate. The test pulse responses before and after the ramp are identical within 2% showing that during this (~3 s) protocol there is no visible outflow of channels into the inactivated state. Panel B shows responses to 1-s steps of different amplitudes with identical test pulses before and after. The trajectories illustrate the MscS closing rates at different tensions, and again, the comparison of test pulse responses shows no inactivation in this time scale. More prolonged steps of 30, 60, 90 and 120 s (shown in panel C) of tension lead to substantial inactivation. Furthermore, the comparison of time courses shows that the equilibrium distribution of the population between C, O and I states is reached within about 90 s, after which the occupancies become time-independent under these conditions (compare with 120 s step).

Based on traces presented in Fig. 2, we picked 120s as the time required for the chain to reach equilibrium. It should be noted that when subjected prolonged steps of tension, patches often become unstable and either rupture or undergo slippage inside the glass pipette, which changes patch position and the radius of curvature. This may introduce errors in probability measurements. To avoid these errors, we employed 1s triangular ramps and recorded the dose response curves throughout the experiment (Fig. 2D) and checked whether the open probability stays within the limits of stochasticity of the experiment and does not drift as a result of changes in membrane curvature.

### Closed↔Open Transitions

As illustrated in Fig. 2A and B, application of quick ramp or pulse protocols causes no inactivation. With the clear separation of time scales for the two competing transitions originating from the closed state, the system subjected to 1-2 s stimuli can be approximated by a two-state model (Kamaraju et al., 2010, 2011; Nakayama et al., 2013). The closed ↔ open branch was probed by a relatively quick 1-s triangular ramp protocol where the tension in the membrane was increased linearly from zero to saturating tension driving all channels to the open state. The response to the symmetric 1-s ramp is shown in Fig 3A, with the ascending (increase of pressure, channel opening) and descending branches (decrease of pressure, channel closure) and a plateau in the middle reflecting response saturation at high pressures.

**Figure 3.**
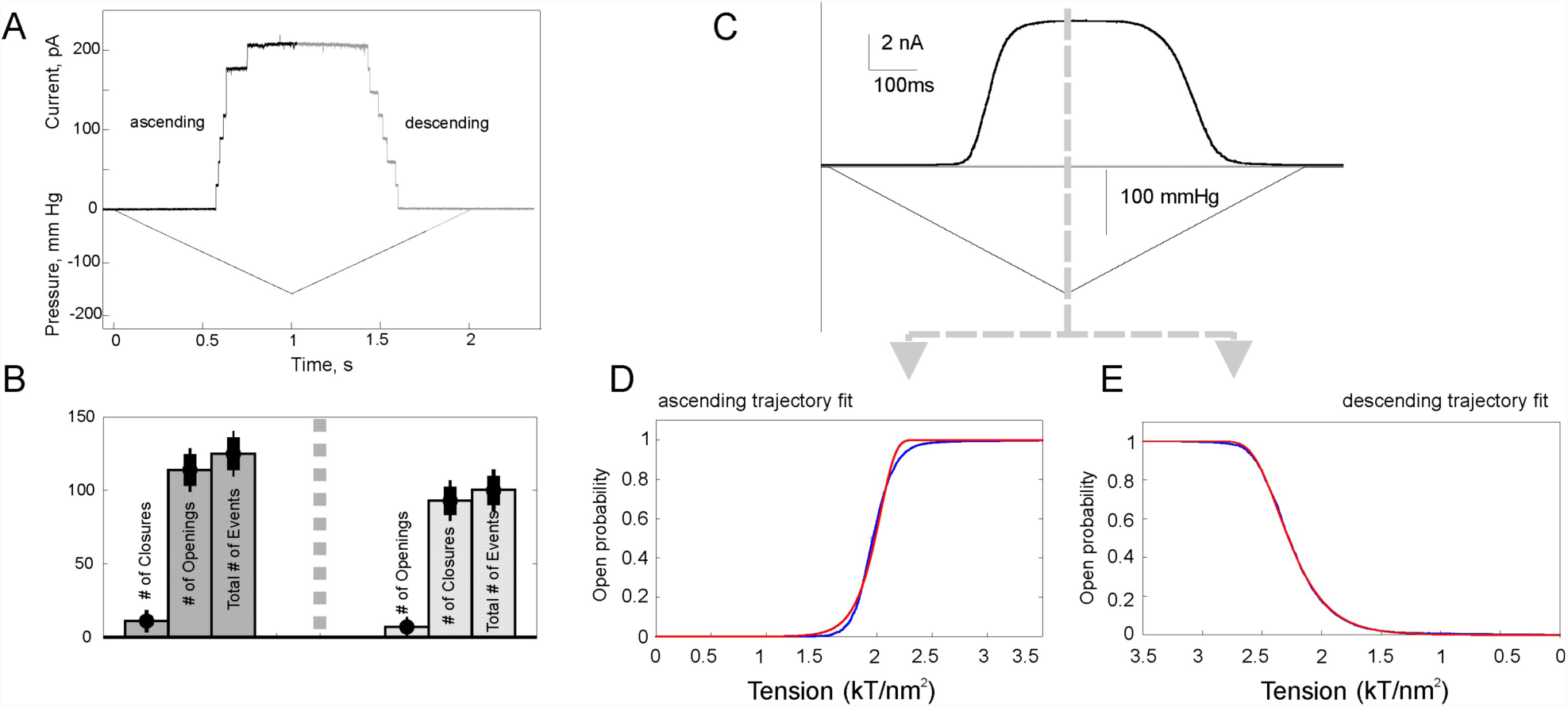
MscS responses to triangular pressure ramps. (A) A ramp response of a small MscS population (8 channels) where individual opening and closing events are seen. (B) Statistics of opening and closing events observed over 20 sequential ramp experiments on several small-population patches. The observed probabilities of closing events during the opening phase and opening events during the closure phase are very low. For this reason, these transitions can be described with a unidirectional kinetics (Eqns. 8 and 9). (C) A response of large (~200 channel) population to a symmetric 1-s triangular ramp. Each leg on the ramp response, ascending (D) and descending (E) represents a separate dose-response curve, which could be fit separately. The probabilities of the open and closed states can be well described by the equations [10, 11] assuming that ***k_0C_*** and ***k_0C_*** are negligible on ascending and descending limbs of the fast ramp protocol respectively (Schlierf et al., 2004). The gating parameters extracted from the fits are listed in Table 1.

**Table 1.** The energy profiles for MscS transitions were tested experimentally and compared with values from literature. For the O ➔ C transition, the area expansion from the open state to the barrier, Δ*A_ob_*, and transition rate in the absence of tension, 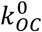, were estimated to be 5.4 *nm*^2^ and 9800 (*s*^−1^) respectively based on the semilog plot of the rates as a function of tension as show in Fig. 4B. These values are in a good agreement with the results obtained by applying the fitting eqn [11] to the descending leg of dose response curve. In the second column, the ΔA values represent the areal distances between the bottoms of the corresponding state wells to the rate-limiting barriers denoted as *b* and *b*^*^, respectively.

| Transition<br>($X \rightarrow Y$) | $\Delta A_{X \rightarrow b, b^*}$<br>($\text{nm}^2$) | Literature value<br>($\text{nm}^2$) | $k_{XY}^0$<br>($\text{s}^{-1}$) | Literature value<br>($\text{s}^{-1}$) |
| --- | --- | --- | --- | --- |
| $C \rightarrow O$ | $6 \pm 1$ | $\sim 11^{(14)}$ | $4\text{e-}6$ | $7\text{e} - 6^{(14)}$ |
| $O \rightarrow C$ | $5 \pm 1$ | $3.8^{(18)}, 4.3^{(3)}, \sim 4.8^{(14)}$ | $1\text{e+}4$ | $2700^{(14)}, 2430^{(22)}$ |
| $C \rightarrow I$ | $5 \pm 1$ | $4.6^{(13)}$ | $2\text{e-}5$ | None |
| $I \rightarrow C$ | $1.2 \pm 0.5$ | $2.6^{(13)}$ | $0.18$ | None |

When tension increases linearly as a fast ramp, the rate of accumulation of open channels can be presented in the form:
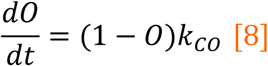

And on the descending leg of dose response curve can be written as:
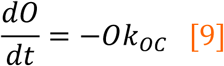

Here we use a simplifying assumption that the rate of closing (*k_oc_*) on the ascending limb and the rate of opening (*k_co_*) on the descending limb are negligible. To verify this assumption, in Fig 3A we provided a typical ramp response of a small channel population where each individual transition is clearly seen. Panel B shows the histogram of the number of closure events compared to the number of openings on the ascending leg of the trace as the tension was quickly increased in a linear fashion. The closing events during tension increase can be considered rare events thus verifying the negligibility of *k_oc_*. Similar reasoning applies to *k_co_* on the descending leg of the dose response. Obviously, only during the quick ramp protocols, the channels make primarily one-way transitions either from the closed to open or from the open to closed state while the membrane tension is linearly increased or decreased with a given rate.

Fig. 3C shows a typical 1-s ramp response of a larger (~200) population of channels which now looks like a smooth curve. To fit the ascending and descending segments of the trace presented in panels D and E we used the solution to the equation [8], which can be written as (Schlierf et al., 2004):
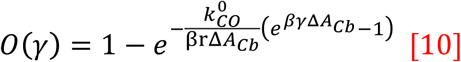

And the closed state probability on the descending leg is given by:
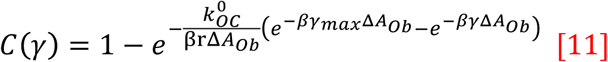

Curves fitting Eqn [10] and Eqn [11] to the ascending and descending legs of the dose response curve (Fig. 3D, E red lines) produced parameters for the closed↔open transition now listed in Table 1.

As a second way of getting the gating parameters for the open à closed transition, we employed the previously described experimental protocol (Kamaraju and Sukharev, 2008; Boer et al., 2011; Nakayama et al., 2013) where the tension is first delivered as a short saturating pulse to pre-condition the entire channel population to the open state and then is changed to various sub-saturating levels. The kinetics of closure is monitored as a function of tension and time. The experiment directly (Fig. 4A) reveals the closing rate, *k_oc_*. Once the logarithm of the rate obtained from the mono-exponential fits of the initial segments of the decaying current traces is plotted against membrane tension, its slope provides information about the Δ*A* between the open well and the rate-limiting barrier, and the y intercept gives the logarithm of the intrinsic closing rate in the absence of tension. This rate might reflect (although may not coincide with) the attempt rate (pre-exponential frequency factor) associated with the transition, as well as the height of the energy barrier. The transition state area (barrier position) and the attempt frequency in the absence of the tension that were captured from the semi-log plot in Fig. 4B were in good agreement with the values extracted from the fit of Eqn [11] to the descending leg of dose response curve as explained in Table 1.

**Figure 4.**
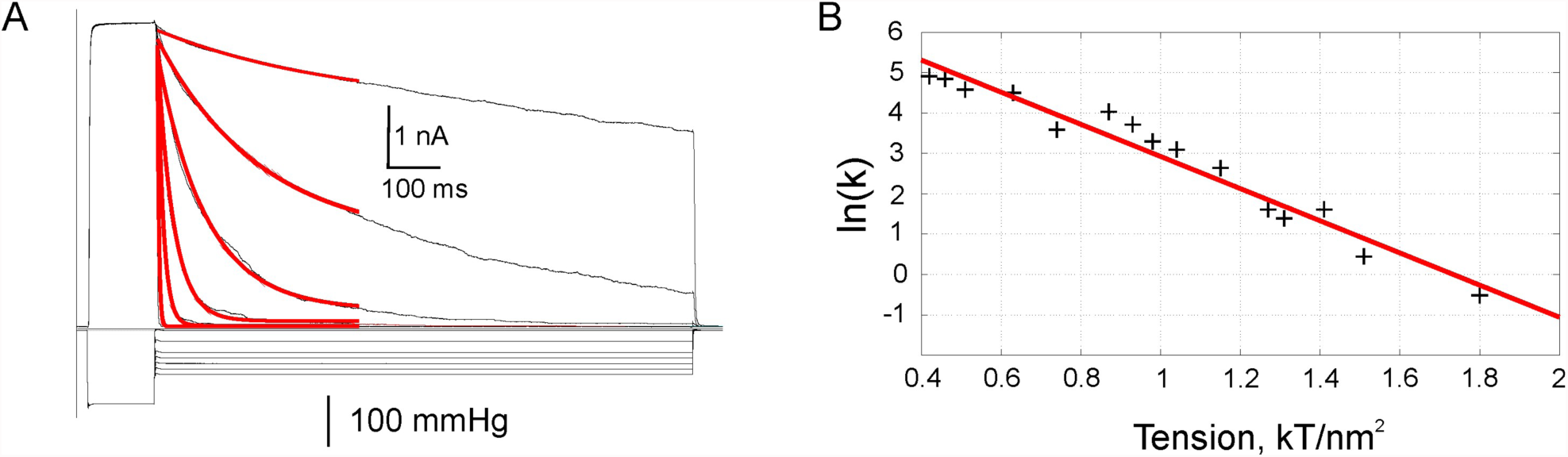
Probing the Open → Closed transition by the pulse-step protocol. (A) The kinetics of the channels pre-conditioned in the open state by the saturating pulse relaxing to the closed state was monitored as a function of tension for 2s and fitted by the mono exponential function to extract the rates. (B) The semi-logarithmic plot of the closing rate as a function of tension. The slope gives an estimate of the areal distance from the bottom of the open-state well to the rate-limiting barrier. The y-intercept suggests the intrinsic closing rate in the absence of tension.

### Closed↔Inactivation Transitions

The gating parameters of the inactivation (C ➔ I) and the reverse (I ➔ C) transitions were recaptured by utilizing multi-step experimental protocols previously described in (Kamaraju and Sukharev, 2008; Kamaraju et al., 2011). The initial short saturating pulse preconditions all the channels in the open state and measures the total number of active channels in the membrane. The tension is then switched to a sub-saturating level for 60s allowing the system to make transitions between the states and to relax toward to the equilibrium distribution determined by tension (Fig. 5A). As the channels made transitions from the open state to the inactivated state while passing through the closed state, namely, O➔C➔I, their distribution was periodically monitored by short saturating pulses interspersed evenly from the beginning to the end enabling us to see the remaining active population. The fraction of the population in each state is deduced using the normalization condition, *P_open_* + *P_Closed_* + *P_inactivated_* = 1. The log of *k_CI_*, which was obtained by fitting the analytic solutions for the model 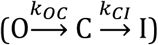 to the experimental traces, was plotted as a function of tension (Fig. 5B). This gives the area from the closed state well to the barrier that separates it from the inactivated state and the logarithm of the intrinsic transition rate in the absence of tension (Table 1).

**Figure 5.**
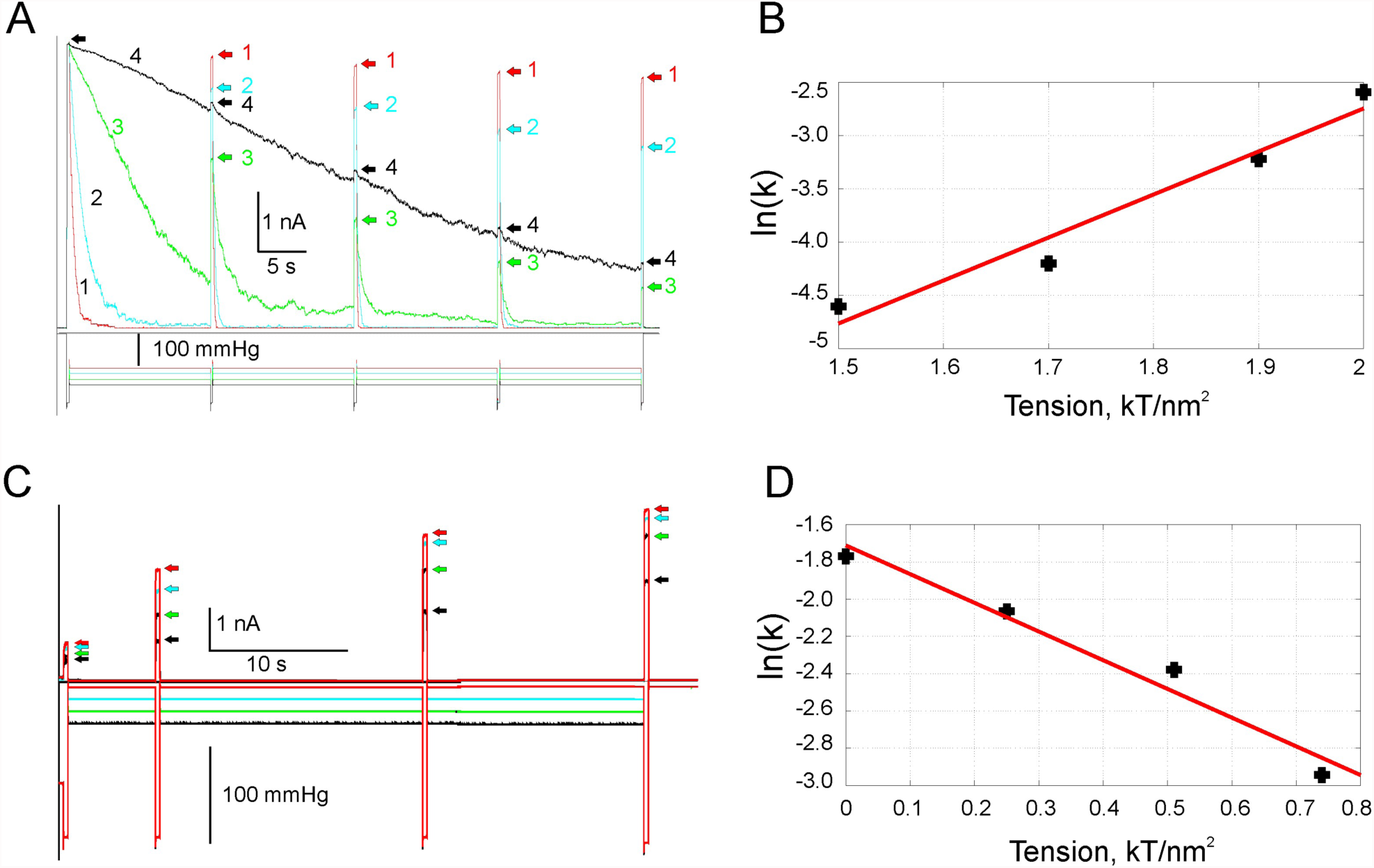
Probing the Closed ↔ Inactivated transition. A) The channels were kept at sub-saturating tensions for 60s, allowing enough time for inactivation to manifest. The number of channels populated in different states was checked by interspersed saturating pulses that demonstrate the distribution of the channels between the states based on the fact that only the channels in the closed state could be activated with tension and the number of the channels in the membrane was constant. With this protocol, as shown in (Kamaraju et al., 2011), channels’ transitions can be described by the following scheme: 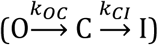 from which rate of the inactivation, ***k_CI_***, was obtained by fitting analytic solutions to experimental data. C) MscS recovery from inactivation is retarded by the membrane tension. Channels were driven to inactivation by a 60-s saturating tension and the extent of recovery from inactivated state was tested by four saturating test pulses applied at different time points for various tension values. The recovery traces were fitted with mono-exponential functions (not shown). (B and D) The semi-log plots of the rates obtained by single exponential fit to data revealed the area expansion and transition rate in the absence of tension associated with the relevant transition. All parameters are listed in Table 1 with experimental uncertainties.

The recovery transition (I ➔ C) was studied by the protocol first reported by (Kamaraju et al., 2011). The channel population is first probed by a short test pulse and then kept at ***γ***\* to maximize inactivation until almost all channels are driven into the inactivated state. After conditioning the channels in the inactivated state at ***γ***\*, tension was decreased to zero or to an intermediate level between 0 and ***γ***\*, and the rate of MscS transition to the closed state was monitored by applying an extended train of short saturating pulses (Fig. 5C). Since the inactivated state cannot open in response to tension, only the channels in the closed state can be activated by tension. Thus, by employing saturating pulses, the recovery transition can be monitored over time. The recovery rate from inactivation slowed down with an increase in the membrane tension as indicated by the red, blue, green and black arrows in Fig 5C, where red arrows show the fastest recovery in the absence of membrane tension. The log of the rates obtained from single exponential fits to data (not shown) was plotted as a function of tension as reported in (Kamaraju et al., 2011)(Fig 5D). The complete set of transition parameters obtained from the slopes and the y intercepts is listed in Table 1.

### Additional Validation of the Model with Experimental Data

Having the kinetic and spatial parameters for the two reversible transitions at hand, we decided to test them in two experiments and compare the results with model prediction. The first experiment approximates a prolonged ‘static’ mechanical perturbation allowing the system to reach equilibrium. The second experiment explores the MscS population in different dynamic regimes under tension ramps applied with different speeds. The first set was designed to compare our analytic results obtained from the steady state solution of the model with the experiments where patches were exposed to 120 s conditioning steps required for the chain to reach equilibrium as shown in Fig. 2C. In Fig. 6, we present the experimental fraction of inactivated channels at the end of a 120 s step shown as blue circles with error bars. The data were obtained from 8 independent stable patches from 3 different spheroplast preparations. The red curve was computed by plugging the parameters listed in Table 1 into *eqn* [4] predicting the steady state inactivation, and the blue curve represents the open probability of the channel on which *γ*_0.5_ is marked with a red circle. Eqn [4 well reproduces the experimentally observed fraction of MscS inactivation as a function of conditioning tension magnitude.

**Figure 6.**
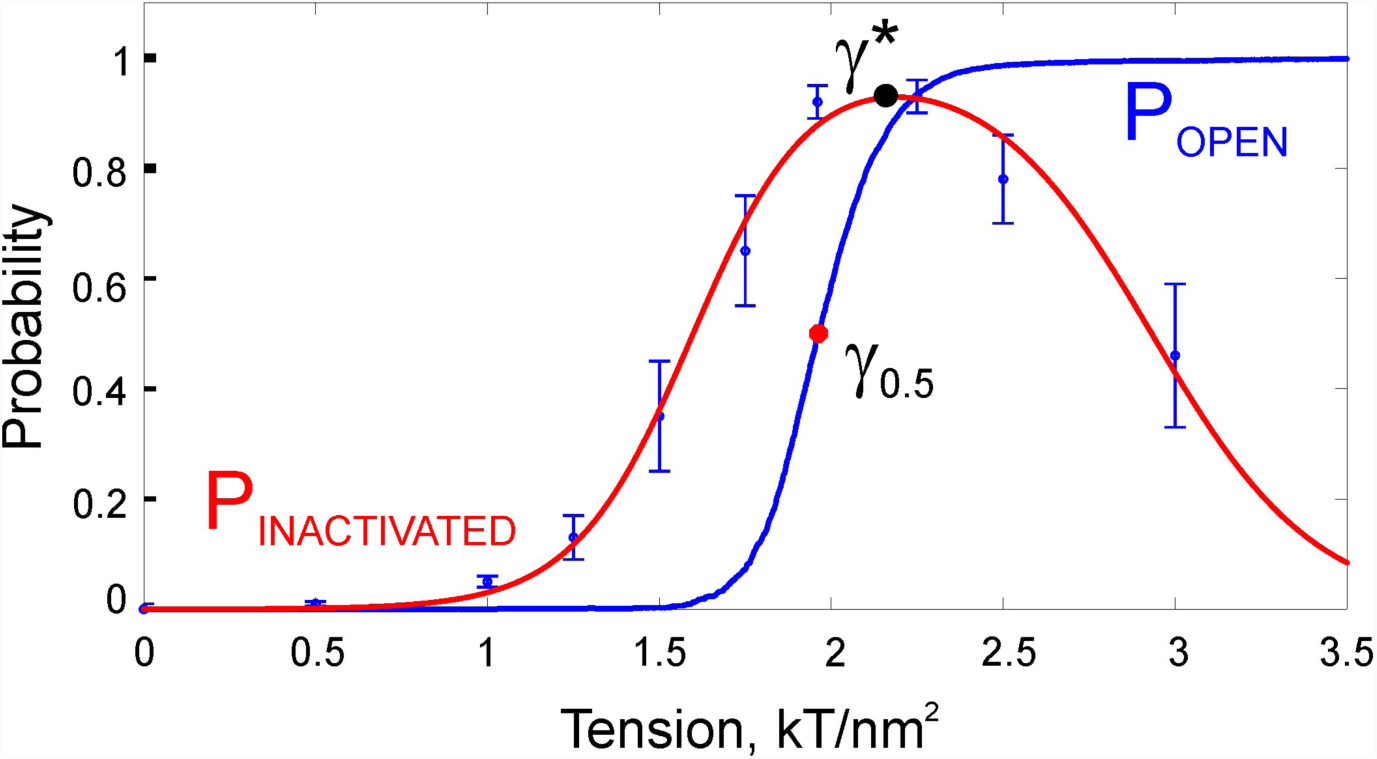
Comparison of experimental steady state (equilibrium) MscS inactivation with 3-state model. The blue data points with the error bars represent steady state MscS inactivation in the course of a 120 s conditioning step as in Fig. 2C, averaged over 8 different membranes. The blue curve is the open probability for the fast (non-equilibrium) activation of channels obtained from a 1s ramp protocol on which, ***γ***_0.5_ is marked as red circle. The red curve, which is the plot of steady state solution of the model **eqn** [4] evaluated with the parameters listed in Table 1, describes the experiments well. The black circle is the tension at which the steady-state inactivation level is maximized, ***γ***\*, predicted by the model, ***eqn*** [6], after plugging in the corresponding values in Table 1.

As previously mentioned, MscS inactivation takes place in a specific range of tensions, and the value of *γ*\* which produces the maximum inactivation is predicted by eqn [6] carrying information about the in-plane area of the inactivated state relative to the area of the open state on the reaction coordinate. The *γ*\* for MscS was calculated to be 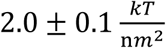 by evaluating eqn [6] with the parameters listed in Table 1 and marked with a black circle on Fig 6, showing a good agreement with the experimental data.

This framework encapsulated in eqn. [7] can be used to determine the inactivated state area for yet unexplored MscS homologs that also show tension-dependent inactivation, but were not subjected to test protocols depicted in Fig 5. In order to illustrate the utility of eqn [7], we calculated the inactivated state area of a newly characterized MscS homolog from *Pseudomonas aeruginosa*, PaMscS-1, which also inactivates with tension (Çetiner et al., 2017). By plugging in the experimental values for the midpoint tension, *γ*_0.5_, and the tension which maximizes the inactivation, *γ*\*, into eqn [7], the inactivated state area of PaMscS-1 was estimated to be 5 ± 1 *nm*^2^, which is comparable with 6 ± 1 *nm*^2^ for EcMscS (see Supplementary Fig. 3). Note that even though a closed-form expression for *γ*\* is obtained from the steady state solution, it is the same tension that maximizes the inactivation at any intermediate time (See Supplementary Fig. 4). The *E. coli* MscS and *P. aeruginosa* MscS-1 share 36 % of identity and likely have the same inactivation mechanism characterized by the same spatial scale of the structural rearrangement, which is presumably the uncoupling of the gate from the lipid-facing helices (Akitake et al., 2007).

Moreover, due to the topology of the transition diagram (Fig. 1), it is also now possible to assign unique energies to the states by choosing a reference state. For example, if we assign the energy of the closed state to be zero, we can uniquely determine the energy of the open and the closed state by using the relation (Schnakenberg, 1976):

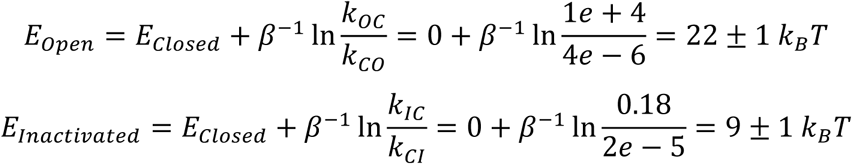

These relationships assign the energies for the bottoms of the open and inactivated state wells on the energy landscape.

To test the correctness of the kinetic constants and their tension dependences obtained in the above experiments, we simulated MscS ramp responses with the coefficients listed in Table 1. In parallel, we employed the same time-dependent ramp protocols where the membrane tension, γ = rt, was linearly raised from zero to its final value of 3 k_B_T/nm^2^ in 1s, 5s, 10s, 30s and 60s, thus changing the rate from fast to slow. As shown previously, MscS channels prefer to fully respond to abrupt stimuli but tend to ignore the slowly applied ones, the behavior that was called the “dashpot” mechanism. Not surprisingly, as the tension increases from zero to the saturating level, it passes through a specific region, which roughly corresponds to γ_0.5_, where the likelihood of switching from the closed to the inactivated state is the highest. At slower rates of stimulus application, channels spend more time in this specific range of tensions and a larger fraction of channels is predicted to end up in the inactivated state. Fig. 7A shows experimental traces (blue) that reflect the fraction of active channels in the patch. All traces were normalized to the amplitude of the shortest ramp response that exhibits the maximal number of active channels. As seen from the bottom part of the graph, tension was raised to its final value of 3 k_B_T/nm^2^ with different rates. Indeed, with the slower rates only about half of the population remains active. The simulation of a 30000-channel Markov chain using the QUBexpress software under the same conditions is shown by the red curves (Fig. 7A, B) which reasonably reproduces the kinetic inactivation mechanism of MscS.

**Figure 7.**
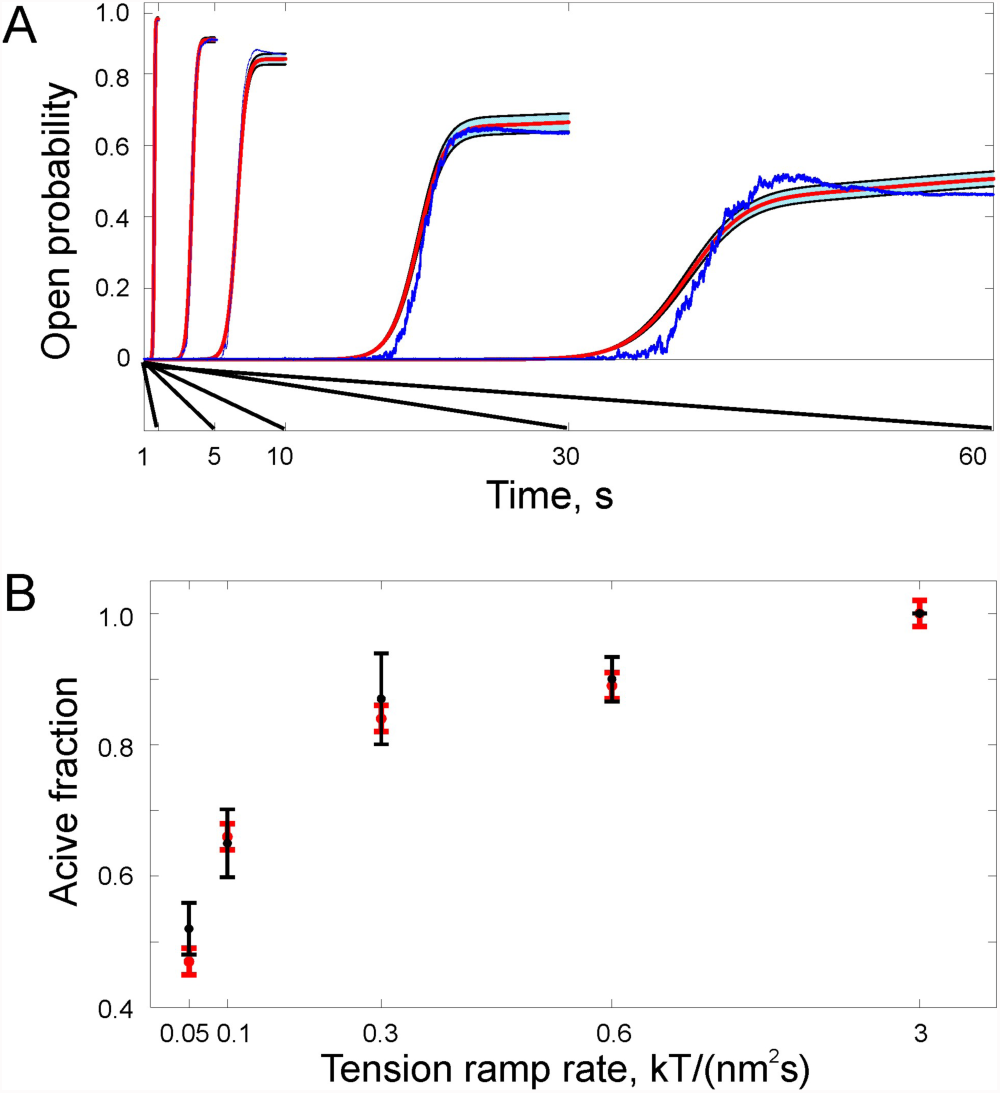
The rate dependency of MscS inactivation. A) The plot of the fraction of the open channels normalized to 1s ramp (fastest). The tension in the membrane was linearly raised to its final value with different rates. At slower rates, channels spent more time in the tension region where inactivation can be achieved. Thus, the slower the application of stimulus, the more likely the channels are to end up in the inactivated state, which can be seen as the decrease in the membrane current. The red curves were the average traces of 30000-channel MscS population simulated by the QUBexpress software with the parameters listed in Table 1 and are in good agreement with the blue traces that correspond to a single realization of the experimental MscS data. B) Relative number of the active channel population with different tension application rates obtained from three independent experiments (black] was successfully described by the model (red).

## Discussion

MS channels in bacteria fulfill the role of osmolyte release valves rescuing cells from mechanical damage in the event of abrupt osmotic downshift (Levina et al., 1999; Boer et al., 2011; Çetiner et al., 2017;). Although the release system in *E. coli* is comprised of MscL, MscS and five other channel species ( Schumann et al., 2010; Edwards et al., 2012;), the low-threshold MscS and the high-threshold MscL were shown to mediate the bulk of osmolyte exchange and each of them is sufficient to rescue the majority of bacterial population from osmotic bursting (Levina et al., 1999). Based on early data, these two channels were deemed partially redundant (Bialecka-Fornal et al., 2015; Booth et al., 2015). Previous comparative studies (Nakayama et al., 2013; Cox et al., 2015) and the analysis of MscS transitions presented above suggest that the channels are specialized. The low-threshold MscS shows characteristic activation and closing properties, and most importantly it inactivates.

The extracted spatiotemporal parameters of MscS transitions (Table 1) predict that the energy landscape for MscS functional cycle must have at least 3 wells for the closed (C), open (O) and inactivated (I) states (Fig. 8A). At zero tension, the opening path predicts a barrier that is located about 55% toward the O well, which is located at a distance of 11 nm^2^ and elevated by about 22 k_B_T relative to the C well. The inactivation profile predicts a smaller distance of about 6.5 nm^2^ between the C and I wells, the energy difference of 9 k_B_T and the rate-limiting barrier located very close (~20 % of the way) to the I well. Application of ‘midpoint’ tension γ_0.5_=7.85 mN/m that tilts the landscape (Fig. 8B) and that puts the bottoms of C and O wells at the same level also drags the I well below the C well prompting massive inactivation.

**Figure 8.**
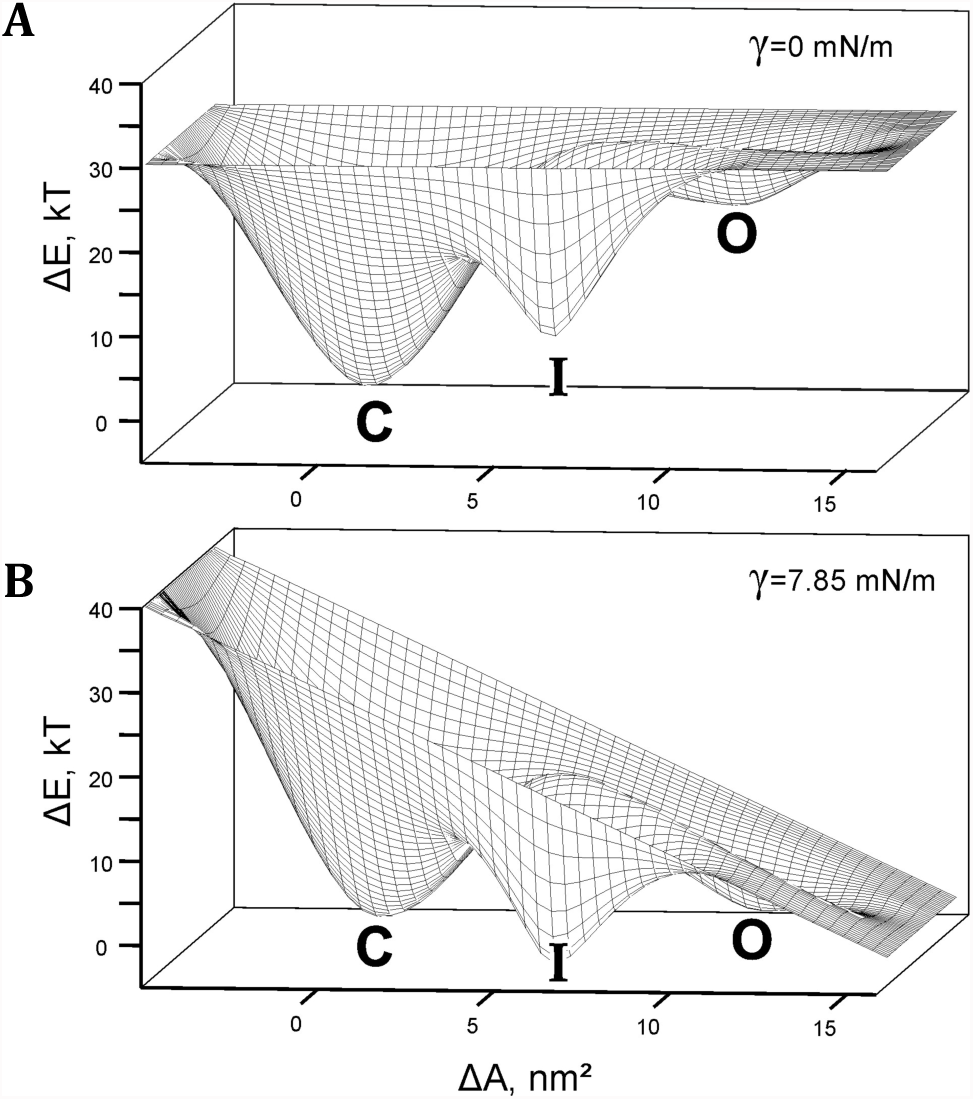
A) The reconstructed energy landscape predicted for MscS in a relaxed membrane (γ=0) and under tension γ_0.5_=7.85 mN/m, at which open and closed states are equally populated (B). In the latter case I state has a lower energy than both the C and O states thus making the inactivated state preferable. The passage to the I state becomes accessible from the C state, whereas a ridge separating O and I states prevents inactivation from the open state. Application of stronger tension skews the landscape more and drives the closed population to the open state. Importantly, inactivated channels at high tensions remain ‘locked’ in the inactivated state.

The gating parameters defined by this profile can be interpreted in the context of its in-vivo function. Subjected to strong (500-1000 mOsm) osmotic downshifts, bacterial cells swell and generate super-threshold tensions in their cytoplasmic membranes within 20-30 ms, and then a release phase with a characteristic time of 30-100 ms begins (Boer et al., 2011; Çetiner et al., 2017). Based on parameters presented in Table 1, when tensions exceed 3 k_B_T/nm^2^ (or 12 mN/m), MscS is predicted to open within 10 ms. The predicted opening rate becomes too fast to be resolved in a typical patch-clamp experiment with existing pressure delivery equipment (HSPC-1), but MscS population will robustly open at sublytic tensions within the swelling time. The opening speed is important because in order to curb swelling the population must be able to release excessive osmolytes faster than the rate of water influx.

Regarding the closing rate, it is estimated to be fast (~10,000 s^-1^) at zero tension. But it is important to keep in mind that as the osmolytes are being dissipated, membrane tension decreases until the channels close. Therefore, channel activity cannot adjust membrane tension below their own activation threshold. The threshold tension thus sets the operating regime for the channel. For MscS, the threshold tension is near 5 mN/m (~1.3 k_B_T/nm^2^) and at this tension the closing rate is predicted to be ~3 s-1, i.e., the channels will be closing in vivo with characteristic time of 0.3 s or slower. This kinetic impediment will provide sufficient time for osmolyte and water exchange between the small cell and the environment in the absence of high intracellular pressure thus reducing tension to a subthreshold level.

As illustrated in Figs. 5, the rates of inactivation and recovery are generally slow compared to the rates of activation and closure, apparently due to a sufficiently high barrier separating the C and I states, smaller expansion during the inactivation, and location of this barrier that is close to the inactivated state. In terms of activation and inactivation, MscS in our experiments essentially functions in two separate time scales. The smaller area change associated with the inactivation process dictates much shallower tension dependence. But what is special about this transition is that with the given parameters inactivation only takes place at moderate tensions above the activation threshold (~1.3 k_B_T/nm^2^) and is significantly boosted at around the activation midpoint (~2.0 k_B_T/nm^2^) where flickering between the open and closed state of the channel is maximized (Fig. 6). It has been analytically shown that the relationship between activation midpoint, *γ*_0.5_ and the tension γ* at which the steady-state degree of activation is the highest depends on the areas of open and inactivated states. This allows estimation of inactivation area without the complex experimental protocols and analysis exemplified in Fig. 5, but rather from a set of more easily accessible parameters (Fig. 6 and supplemental Fig. 3). The inactivation area in turn may help interpreting structural models of inactivation (Anishkin and Sukharev, 2004; Vásquez et al., 2008). The reconstructed landscape well predicts ‘smart’ behavior of fully responding to abrupt stimuli (emergency situations) but ignoring tension stimuli which are applied slowly (non-emergency situations).

The specific role of the low-threshold inactivating MscS in the osmotic response and functional cooperation with the non-inactivating (two-state) high-threshold MscL channel may be envisioned in the following way. At low-magnitude shock, the low-threshold MscS, may completely fulfill the pressure/volume adjustment without engaging MscL. At higher shocks, MscL will activate and take the major part in the fast osmolyte release, but when tension drops down to MscL threshold (~9 mN/m), it will close and the tension adjustment will be stalled at that level. Under these conditions, the non-inactivating MscL will still be able to flicker to the open state, which will be disruptive for vital gradients and cell energetics. MscS appears to be a critical asset in this situation. Fully open at 9 mN/m (~2.3 k_B_T/nm^2^), MscS will continue the dissipation process to take membrane tension considerably below MscL threshold. At its own activation midpoint of 7.8 mN/m (~2 k_B_T/nm^2^) or below MscS will inactivate and this would be a proper leak-free termination of the osmotic permeability response. This picture should be augmented by the findings that the rate of inactivation (but not the preferable tension range) sharply increases in the presence of cytoplasmic crowders (Grajkowski et al., 2005; Rowe et al., 2014). The increased macromolecular excluded volume is an indicator of the cytoplasm “over-draining” and the signal for faster MscS inactivation that prevents small osmolyte and water extrusion.

We conclude that the existence of a non-conductive, tension-insensitive (inactivated) state and the location of the inactivated state well on the energy landscape relative to other states are not coincidental but rather a result of a billion-year evolution to provide a more efficient and ‘economic’ response to osmotic challenges thus contributing to the osmotic fitness of bacteria in the ever changing environment.

## ACKNOWLEDGMENTS

The work was supported by NIH R21AI105655 and GM107652 grants to SS. UC was supported by the U.S. Department of Education GAANN ‘Mathematics in Biology’ Fellowship. UC is also indebted to Drs. Andriy Anishkin and Oren Raz (University of Maryland, College Park) and Yigit Subaçi (Los Alamos National Laboratory) for their stimulating discussions. The authors thank Ms. Stephanie Sansbury for cloning MscS into tightly-regulated pBAD expression system and Madolyn Britt for editorial comments.

## Bibliography

Akitake, B., Anishkin, A., and Sukharev, S. (2005). The “dashpot” mechanism of stretch-dependent gating in MscS. J Gen Physiol 125, 143-154.

Akitake, B., Anishkin, A., Liu, N., and Sukharev, S. (2007). Straightening and sequential buckling of the pore-lining helices define the gating cycle of MscS. Nat Struct Mol Biol 14, 1141-1149.

Alloui, A., Zimmermann, K., Mamet, J., Duprat, F., Noël, J., Chemin, J., Guy, N., Blondeau, N., Voilley, N., Rubat-Coudert, C., et al. (2006). TREK-1, a K+ channel involved in polymodal pain perception. EMBO J 25, 2368-2376.

Anishkin, A., and Sukharev, S. (2004). Water dynamics and dewetting transitions in the small mechanosensitive channel MscS. Biophys J 86, 2883-2895.

Anishkin, A., Kamaraju, K., and Sukharev, S. (2008a). Mechanosensitive channel MscS in the open state: modeling of the transition, explicit simulations, and experimental measurements of conductance. J Gen Physiol 132, 67-83.

Anishkin, A., Akitake, B., and Sukharev, S. (2008b). Characterization of the resting MscS: modeling and analysis of the closed bacterial mechanosensitive channel of small conductance. Biophys J 94, 1252-1266.

Bell, G.I. (1978). Models for the specific adhesion of cells to cells. Science 200, 618-627.

Belyy, V., Kamaraju, K., Akitake, B., Anishkin, A., and Sukharev, S. (2010a). Adaptive behavior of bacterial mechanosensitive channels is coupled to membrane mechanics. J Gen Physiol 135, 641-652.

Belyy, V., Anishkin, A., Kamaraju, K., Liu, N., and Sukharev, S. (2010b). The tension-transmitting “clutch” in the mechanosensitive channel MscS. Nat Struct Mol Biol 17, 451-458.

Bialecka-Fornal, M., Lee, H.J., and Phillips, R. (2015). The rate of osmotic downshock determines the survival probability of bacterial mechanosensitive channel mutants. J Bacteriol 197, 231-237.

Blount, P., Sukharev, S.I., Moe, P.C., Schroeder, M.J., Guy, H.R., and Kung, C. (1996). Membrane topology and multimeric structure of a mechanosensitive channel protein of Escherichia coli. EMBO J 15, 4798-4805.

Boer, M., Anishkin, A., and Sukharev, S. (2011). Adaptive MscS gating in the osmotic permeability response in E. coli: the question of time. Biochemistry 50, 4087-4096.

Booth, I.R., Miller, S., Müller, A., and Lehtovirta-Morley, L. (2015). The evolution of bacterial mechanosensitive channels. Cell Calcium 57, 140-150.

Çetiner, U., Rowe, I., Schams, A., Mayhew, C., Rubin, D., Anishkin, A., and Sukharev, S. (2017). Tension-activated channels in the mechanism of osmotic fitness in Pseudomonas aeruginosa. J Gen Physiol 149, 595-609.

Cox, C.D., Nakayama, Y., Nomura, T., and Martinac, B. (2015). The evolutionary “tinkering” of MscS-like channels: generation of structural and functional diversity. Pflugers Arch 467, 3-13.

Edwards, M.D., Black, S., Rasmussen, T., Rasmussen, A., Stokes, N.R., Stephen, T.L., Miller, S., and Booth, I.R. (2012). Characterization of three novel mechanosensitive channel activities in Escherichia coli. Channels (Austin) 6, 272-281.

Grajkowski, W., Kubalski, A., and Koprowski, P. (2005). Surface changes of the mechanosensitive channel MscS upon its activation, inactivation, and closing. Biophys J 88, 3050-3059.

Haswell, E.S., Peyronnet, R., Barbier-Brygoo, H., Meyerowitz, E.M., and Frachisse, J.M. (2008). Two MscS homologs provide mechanosensitive channel activities in the Arabidopsis root. Curr Biol 18, 730-734.

Heurteaux, C., Lucas, G., Guy, N., El Yacoubi, M., Thümmler, S., Peng, X.D., Noble, F., Blondeau, N., Widmann, C., Borsotto, M., et al. (2006). Deletion of the background potassium channel TREK-1 results in a depression-resistant phenotype. Nat Neurosci 9, 1134-1141.

Kamaraju, K., and Sukharev, S. (2008). The membrane lateral pressure-perturbing capacity of parabens and their effects on the mechanosensitive channel directly correlate with hydrophobicity. Biochemistry 47, 10540-10550.

Kamaraju, K., Gottlieb, P.A., Sachs, F., and Sukharev, S. (2010). Effects of GsMTx4 on bacterial mechanosensitive channels in inside-out patches from giant spheroplasts. Biophys J 99, 2870-2878.

Kamaraju, K., Belyy, V., Rowe, I., Anishkin, A., and Sukharev, S. (2011). The pathway and spatial scale for MscS inactivation. J Gen Physiol 138, 49-57.

Kung, C., Martinac, B., and Sukharev, S. (2010). Mechanosensitive channels in microbes. Annu Rev Microbiol 64, 313-329.

Levina, N., Tötemeyer, S., Stokes, N.R., Louis, P., Jones, M.A., and Booth, I.R. (1999). Protection of Escherichia coli cells against extreme turgor by activation of MscS and MscL mechanosensitive channels: identification of genes required for MscS activity. EMBO J 18, 1730-1737.

Martinac, B., Buechner, M., Delcour, A.H., Adler, J., and Kung, C. (1987). Pressure-sensitive ion channel in Escherichia coli. Proc Natl Acad Sci U S A 84, 2297-2301.

Nakagawa, Y., Katagiri, T., Shinozaki, K., Qi, Z., Tatsumi, H., Furuichi, T., Kishigami, A., Sokabe, M., Kojima, I., Sato, S., et al. (2007). Arabidopsis plasma membrane protein crucial for Ca2+ influx and touch sensing in roots. Proc Natl Acad Sci U S A 104, 3639-3644.

Nakayama, Y., Yoshimura, K., and Iida, H. (2013). Electrophysiological characterization of the mechanosensitive channel MscCG in Corynebacterium glutamicum. Biophys J 105, 1366-1375.

Norris, J.R. (1998). Markov chains (Cambridge university press).

Okada, K., Moe, P.C., and Blount, P. (2002). Functional design of bacterial mechanosensitive channels. Comparisons and contrasts illuminated by random mutagenesis. J Biol Chem 277, 27682-27688.

Prole, D.L., and Taylor, C.W. (2013). Identification and analysis of putative homologues of mechanosensitive channels in pathogenic protozoa. PLoS ONE 8, e66068.

Retailleau, K., and Duprat, F. (2014). Polycystins and partners: proposed role in mechanosensitivity. J Physiol (Lond) 592, 2453-2471.

Ritort, F. (2004). Work Fluctuations, Transient Violations of the Second Law and Free-Energy Recovery Methods: Perspectives in Theory and Experiments. In Poincaré Seminar 2003, J. Dalibard, B. Duplantier, and V. Rivasseau, eds. (Basel: Birkhäuser Basel), pp. 193-226.

Rowe, I., Anishkin, A., Kamaraju, K., Yoshimura, K., and Sukharev, S. (2014). The cytoplasmic cage domain of the mechanosensitive channel MscS is a sensor of macromolecular crowding. J Gen Physiol 143, 543-557.

Schlierf, M., Li, H., and Fernandez, J.M. (2004). The unfolding kinetics of ubiquitin captured with single-molecule force-clamp techniques. Proc Natl Acad Sci U S A 101, 7299-7304.

Schnakenberg, J. (1976). Network theory of microscopic and macroscopic behavior of master equation systems. Rev. Mod. Phys. 48, 571-585.

Schumann, U., Edwards, M.D., Rasmussen, T., Bartlett, W., van West, P., and Booth, I.R. (2010). YbdG in Escherichia coli is a threshold-setting mechanosensitive channel with MscM activity. Proc Natl Acad Sci U S A 107, 12664-12669.

Sotomayor, M., and Schulten, K. (2004). Molecular dynamics study of gating in the mechanosensitive channel of small conductance MscS. Biophys J 87, 3050-3065.

Sotomayor, M., van der Straaten, T.A., Ravaioli, U., and Schulten, K. (2006). Electrostatic properties of the mechanosensitive channel of small conductance MscS. Biophys J 90, 3496-3510.

Steinfeld, J.I., Francisco, J.S., and Hase, W.L. (1989). Chemical kinetics and dynamics (New Jersey: Prentice Hall Englewood Cliffs).

Sukharev, S.I., Martinac, B., Arshavsky, V.Y., and Kung, C. (1993). Two types of mechanosensitive channels in the Escherichia coli cell envelope: solubilization and functional reconstitution. Biophys J 65, 177-183.

Sukharev, S.I., Sigurdson, W.J., Kung, C., and Sachs, F. (1999). Energetic and spatial parameters for gating of the bacterial large conductance mechanosensitive channel, MscL. J Gen Physiol 113, 525-540.

Vásquez, V., Sotomayor, M., Cordero-Morales, J., Schulten, K., and Perozo, E. (2008). A structural mechanism for MscS gating in lipid bilayers. Science 321, 1210-1214.

Volkers, L., Mechioukhi, Y., and Coste, B. (2015). Piezo channels: from structure to function. Pflugers Arch 467, 95-99.

Walsh, C.M., Bautista, D.M., and Lumpkin, E.A. (2015). Mammalian touch catches up. Curr Opin Neurobiol 34, 133-139.

